# Lexical and Semantic Coding in the Occipital Cortex of Sighted and Blind

**DOI:** 10.64898/2026.07.22.739989

**Authors:** Giuliano Giari, Lorenzo Vignali, Yangwen Xu, Olivier Collignon, Davide Crepaldi, Roberto Bottini

## Abstract

How early visual cortex (EVC) supports language and semantic processing in sighted and congenitally blind individuals remains debated. Some predict that blindness induces radical functional reorganization of EVC, whereas others suggest more modest scaling-up changes of a neurofunctional architecture present in sighted individuals. We recorded whole-head MEG in 19 early blind (EB) and 21 sighted adults (SC) listening to spoken adjectives belonging to different semantic categories (i.e., abstract, concrete visual, concrete non-visual) and performing lexical and semantic decision tasks. In the lexical task, lexical information (i.e., word vs pseudoword) could be reliably decoded in perisylvian language areas and the EVC of both groups, with group differences (EB > SC) localized to occipital areas between 0.4 and 0.8 s after word onset. During the semantic task, semantic category information could be decoded in a network overlapping with the canonical semantic system and encompassing the EVC of both groups, with group differences (EB > SC) localized to occipital areas between 0.9 and 1 s after stimulus onset. We further characterized EVC properties and showed that i) EVC semantic information decoding is task sensitive, with no above-chance decoding in EB or SC in the lexical task ii) in the blind EVC it was possible to decode abstract from concrete concepts but not visual from non-visual concepts, a profile similar to that of posterior cingulate areas, but different to anterior cingulate cortex which showed above chance classification for all semantic categories. These results suggest that deprived EVC is integrated into a distributed lexical–semantic network carrying behaviourally relevant semantic information, although to a different extent.

## Introduction

Understanding how the visual brain functionally reorganizes in individuals born without sight is a fundamental question in cognitive neuroscience. When deprived of visual input, are these regions left functionally dormant, or do they instead support new sensory and cognitive functions? Crossmodal plasticity refers to the brain’s ability to reorganize such that one sensory modality can compensate for the loss of another (Bavelier and Neville, 2002). In congenital blindness, crossmodal plasticity is robustly documented: “visual” occipital cortices become attuned to other modalities, including auditory and haptic/tactile perception (Voss, 2019; Collignon et al., 2009, 2011; Heimler et al., 2014; Burton, 2002, 2003, 2006; Matuszewski et al., 2025). Most strikingly, however, blind “visual” areas are not only recruited by other senses, but also participate in higher-cognitive processes such as language and mathematics (Bedny et al., 2011; Crollen et al., 2019). Visual cortices, including primary visual areas, process linguistic information in many ways comparable to classic perisylvian language regions: they show sensitivity to intelligibility (Van Ackeren et al., 2018), semantics (Röder et al., 2002; Noppeney et al., 2003), syntactic complexity (Röder et al., 2002; Lane et al., 2015), linguistic categories such as nouns and verbs (Urbaniak et al., 2025), complex combinatorial structure (Bedny et al., 2011), and tasks involving verbal memory and verb generation (Amedi et al., 2003).

Evidence for higher cognitive functions and, in particular, language processing led some authors to put forward a *cognitive pluripotency* hypothesis of the blind occipital areas (Bedny, 2017). According to this view, the human cortex – including “visual” occipital cortex – is not innately hard-wired for a single sensory domain, but is a flexible computational resource that can, in principle, implement a wide range of cognitive functions. Which function a given patch of cortex comes to support is determined by the pattern of inputs it receives during development and by its long-range connectivity, rather than by a genetically determined blueprint (Bedny, 2017; Kanjlia et al., 2016). In congenital blindness, the absence of visual input frees occipital circuitry to be recruited by higher cognitive systems such as language, numerical cognition, and executive control.

However, the cognitive pluripotency hypothesis has not gone unchallenged, as several objections have been raised against a pluripotent role of visual areas in congenitally blind individuals (Makin & Krakauer, 2023; Fine & Park, 2018). Most notably, Makin argues for insufficient evidence of dramatic functional reorganization in the brain after sensory deprivation. For instance, the categorical organization of the ventral occipito-temporal cortices seems spared by blindness (Mattioni et al., 2020) as well as by transient periods of visual deprivation (Mattioni et al., 2025). From this perspective, changes in function after injury or deprivation can be better explained by plasticity within pre-existing architectures, a case of remapping or unmasking of latent inputs (Amedi et al., 2003, Ricciardi & Pietrini, 2011, Collignon et al., 2013). With respect to language processing in the occipital cortex of blind individuals, the functional remapping hypothesis implies that language-related responses are not a newly acquired function, but should already be present in the occipital cortex of sighted participants. This conclusion found preliminary validation in recent studies investigating language processing in primary visual areas of sighted adults (Seydell-Greenwald et al., 2023; Venezia et al., 2021). However, these results await further validation, as studies including both sighted and blind participants, and focused on activation differences (i.e., blind > sighted or sighted > blind) did not find sighted-specific activations that would reveal the latent precursors of nonvisual functions in sighted people’s visual cortex (Burton et al., 2002, 2003; Bedny et al., 2011; Lane et al., 2015; Amedi et al., 2003; Röder et al., 2000).

Although language sensitivity in the occipital cortex of congenitally blind individuals is now well documented—during Braille reading, verb generation, sentence comprehension, and verbal memory (e.g., Amedi et al., 2003, 2004; Bedny et al., 2012; Burton et al., 2002, 2003; Sadato et al., 1996)—there is comparatively little direct evidence that processing in early visual cortex (EVC) goes beyond lexical or phonological operations to encode genuinely semantic representations. Suggestions that this may be the case come from a small number of recent findings. First, multivoxel decoding studies using *non-linguistic* natural sounds show that early visual areas (V1–V3) in congenitally blind individuals and blindfolded sighted controls carry information about semantic *sound* categories (e.g., humans, animals, vehicles, tools/objects), over and above low-level acoustic structure (Vetter, Smith, & Muckli, 2014; Pollicina, Müller, Dalton, & Vetter, 2025; Mattioni et al., 2020). In these studies, category-specific patterns in EVC generalize across different exemplars within a category, suggesting that EVC can encode abstract categorical properties of sounds, even though the stimuli are not linguistic. Moreover, neurostimulation work reports that disrupting the occipital pole with repetitive TMS selectively impairs verb generation, with predominantly *semantic* (rather than phonological or articulatory) errors (Amedi et al., 2004). Taken together, these strands of evidence hint that, at least under some conditions, occipital cortex—and possibly EVC—can participate in semantic-level processing, but the evidence is sparse, leaving open how deep and systematic semantic representations in early “visual” cortex truly are.

Thus, it remains unclear whether, and under which specific conditions, linguistic—and in particular semantic—information is processed in the early visual cortex of congenitally blind individuals, and to what extent such processing differs from that observed in sighted people. Most of the available evidence comes from fMRI studies and therefore provides a temporally coarse picture. As a result, virtually nothing is known about the **time course** of language processing in these regions in either group: we do not know when EVC becomes engaged during comprehension, which stages of lexical–semantic processing it tracks, or how its temporal profile compares to that of classical language and semantic regions in the brain.

To address these questions, we recorded MEG signals in a group of early blind individuals and a group of sighted controls while they listened to single spoken adjectives. The adjectives belonged to three semantic categories: strictly visual (e.g., *bright, red*), non-visual but concrete (e.g., *loud, salty*), and abstract (e.g., *mental, confused*). Across two separate sessions, with the very same stimuli, participants performed either a lexical decision task (discriminating words from phonotactically plausible nonwords) or a semantic decision task (identifying words that referred to sensory experiences versus words that did not). This design allowed us to test whether early visual cortex in early blind individuals encodes both lexical and semantic properties of auditory words, whether similar coding is present in sighted participants, and to characterize the time course of this processing in both groups. Critically, by comparing lexical and semantic tasks with identical auditory input, we could also ask whether semantic processing in the deprived visual cortex is task-dependent.

## Materials & Methods

### Participants

A total of forty participants took part in the experiment. Nineteen participants were early blind (EB; 9 females, Age M=34±7SD years). All EB participants lost their sight either at birth or within the first 3 years of life, reporting faint light perception and no visual memories. Twenty-one participants were recruited as the sighted control group (SC; 9 females; Age M=36±7.6SD years). Groups were matched by age and years of education (EB=15.4±2.8, SC=15.9±2.2). All participants were Italian native speakers and reported no history of neurological or psychological disorders. The experiment was approved by the Ethical committee of the University of Trento and was carried out in accordance with the Declaration of Helsinki. Participants gave written informed consent before starting the experiment and were reimbursed for their participation.

### Stimuli

Thirtynine Abstract (A, e.g., *confuso*/confused, *mentale*/mental), 39 unimodal-visual (V, e.g., *rosso*/red, *sbiadito*/faded) and 39 concrete-non-visual (C, e.g., *ruvido*/rough; *saporito*/tasty) words were selected from an italian database of modality exclusivity norms (Morucci et al., 2019). Following Connell and Lynott (2012) we used max perceptual strength (MPS) - a measure of the dominant perceptual modality- to distinguish between abstract and concrete words. The list of abstract words includes items with low values of MPS (M=2.98, SD = .58), whereas unimodal visual (M = 4.9, SD = .12) and concrete-non-visual (M = 4.8, SD = .29) words were very high in MPS. In order to distinguish V words from C and A words, we relied on the visual mean score, showing higher values for V (M = 4.8, SD = .16) as compared to C (M = 2.57, SD = .82) and A (M = 2.81, SD = .55). Overall values for frequency, number of syllables, orthographic neighbours, age of acquisition, first syllable frequency, familiarity score and contextual availability were matched across the three classes (see Table 1).

**Table 1.** Linguistic variables across the three semantic categories. Zipf: linguistic frequency expressed as zipf value; NSyll: number of syllables; ON: orthographic neighbours; AOA: age of acquisition, expressed in years of age; FSyll: first syllable frequency; Fam: familiarity; CA: contextual availability; Values indicate mean score and standard deviation in brackets.

| Class | Zipf | NSyll | ON | AOA | FSyll | Fam | CA |
| --- | --- | --- | --- | --- | --- | --- | --- |
| A | 3.2(.93) | 3.3(.77) | 8.33(11.2) | 9.56(2.3) | 8.01(2.06) | 5.71(.95) | 4.97(.83) |
| C | 2.85(.84) | 3.25(.59) | 6.46(7.77) | 7.6(1.94) | 7.43(2.5) | 5.45(1.01) | 5.86(.83) |
| V | 2.97(.94) | 3.18(.82) | 7.5(11.2) | 7.2(2.6) | 6.93(2.89) | 5.39(1.23) | 5.91(.98) |

For the lexical decision task, we created pseudowords by submitting the original stimulus set (i.e., 117 words) to the software Wuggy (Keuleers & Brysbaert, 2010). The resulting pseudowords were automatically matched (pairwise) to original stimuli for number of syllables, word length, number of letters, and stress pattern. However, with a total of 234 stimuli, the lexical decision task would have lasted double as much as the semantic task. Partial balancing of tasks durations was achieved by the inclusion of an additional 71 word-fillers to the semantic task. These were matched for word frequency and number of syllables to the original set and will not be considered for further analysis. All stimuli were then synthesized with an artificial female voice (Talk-ToMe software) and recorded using Audio Hijack Pro in a single wave file including all the lexical items (recording format: 16 bits, sample frequency of 44.1 kHz). Individual items were then selected and saved as separate wav files. The amplitude of each audio file was normalized, and the silence before and after the spoken word was cut using the software Praat (Boersma et al., 2018). Finally, the duration of each audio file was adjusted to the number of syllables, that is: two-syllable words lasted 0.4 s, three-syllable words 0.6 s and four-syllables words 0.8 s.

### Experimental Design

On the same day, participants performed the lexical decision task and the semantic task in two recording sessions, separated by a long (∼30min) break, with the session order counterbalanced across participants. During the lexical decision task (LEX) participants were instructed to discriminate between words and pseudowords (i.e., is the word you heard a real word or a meaningless sound?). During the semantic task (SEM), participants were instructed to discriminate between abstract and concrete words (i.e., is the word you heard related to the senses?). Crucially, the same word stimuli were used in both tasks, ensuring that all differences in processing reflect the influence of the task instructions alone. Participants were familiarized with a short version of the task (30 trials taken from a different stimulus set) on a portable PC outside the MEG chamber and were blindfolded before entering the MEG.

Stimulus presentation was controlled via MATLAB and Psychtoolbox (Brainard 1997) and were delivered - at a comfortable sound intensity - via loudspeakers in the MEG chamber. Each stimulus presentation was preceded by a 1.5 s silence interval and followed by a 2 s period. A short sound (i.e., a beep) then initiated the response interval, lasting an additional 2 s, after which there was an additional inter-trial silence interval jittered between 0.5 and 1 s. Responses were recorded using an MEG-compatible button box (Vpixx Technologies, Canada) operated with the dominant hand’s index and middle fingers. Crucially, we took several precautions to minimize potential confounds between motor execution and cognitive processes of interest. First, having a delayed period for the response ensured a greater temporal dissociation between word processing and motor preparation and execution. Second, because during each session individual stimuli appeared twice, the response mapping was counterbalanced between occurrences. To illustrate, if words were initially detected via right-button press and pseudowords via left-button press, in the second half of the session, the buttons (and the instructions) were inverted, words = left and pseudowords = right. Each testing session lasted approximately 2 h and was divided into six seven-minute runs separated by five short breaks.

### MEG data acquisition and preprocessing

MEG signals were recorded at a sampling rate of 1 kHz using a whole-head 306 sensor (204 planar gradiometers; 102 magnetometers) Vectorview system (Elekta Neuromag, Helsinki, Finland) placed in a magnetically shielded room (AK3B, Vakuumschmelze, Hanau, Germany) at the Center for Mind/Brain Sciences of the University of Trento. The MEG signal was band- pass filtered online between 0.1 and 330 Hz. To continuously determine the position of head in relation to the MEG helmet, five head-position indicator coils (HPIs) were employed. Prior to the recording, individual head shapes were digitized (minimum 300 points) along with fiducial points (the nasion and the left and right pre-auricular points) using a Polhemus FASTRAK 3D digitizer (Fastrak Polhemus, Inc., Colchester, VA, USA). A spatio-temporal variant of signal-space separation (SSS) algorithm implemented in the MaxFilter 2.0 software (Elekta Neuromag®) was used to isolate external sources of noise from the head-generated signals. During this procedure, bad channels identified through visual inspection (<12 channels per run) were replaced by interpolation. Finally, data from the different sessions were aligned to an individual common head position (MaxMove). Data analysis was done using non-commercial software packages such as Fieldtrip (Oostenveld, Fries, Maris, & Schoffelen, 2011), MNE-Python (Gramfort et al., 2013) and custom scripts in MATLAB 2019a and Python (3.10). Continuous MEG recordings were high-pass filtered using a two-pass Butterworth filter with a cutoff frequency of 0.1 Hz and segmented from -1.5 s before to 2 s after stimulus onset. Epochs were then downsampled to 250 Hz and time segments contaminated by artifacts were manually rejected using the function *ft_rejectvisual*. Epochs that were above 2 in ‘z-value’ and above 14 of ‘kurtosis’ were excluded from further analyses (total data loss of M = 6%, SD = 4%). Clean epochs underwent Independent Component Analysis (ICA) with the ’runica’ algorithm. The topographies and time series of the 30 components explaining the most variance were visually inspected to identify and remove those that matched the stereotypical cardiac and ocular artifacts. Remaining components were back-projected to the original sensor-space and low-pass filtered with a 40 Hz Butterworth filter before baseline correction was applied with respect to a -0.5 to -0.1 s pre-stimulus time window.

### Source reconstruction

Distributed minimum-norm source estimation (Hämäläinen & Ilmoniemi, 1994) was applied following the standard procedure in MNE-Python (Gramfort et al., 2013). Anatomical T1- weighted MRI images were acquired during a separate session in a MAGNETOM Prisma 3 T scanner (Siemens, Erlangen, Germany) using a 3D MPRAGE sequence, 1mm3 resolution, TR = 2140 ms, TI = 900 ms, TE = 2.9 ms, flip angle 12°. Anatomical MRI images were processed using an automated segmentation algorithm of the Freesurfer software (Fischl, 2012). Co-registration of MEG sensor configuration and the reconstructed scalp surfaces was based on digitized fiducial points matched on anatomical locations on structural images and ∼300 head shape points. Four sighted participants did not undergo the MRI session, this procedure was thus used to morph the Freesurfer template to match their digitised head shape. We then defined a single shell boundary element model and the white matter boundary was used to define a surface source space with ‘oct6’ spacing, resulting in 4096 vertices per hemisphere. The noise covariance matrix was calculated from the prestimulus interval (-0.2s to 0s) of the different trials, reducing its rank to the degrees of freedom left after application of Maxfilter (Westner et al., 2022). The inverse solution was regularised with the default lambda value of 0.1 and restricted to the orientation normal to the cortical surface. The inversion matrix was then used to project single-trial time courses to the source level.

### Classification analysis

Classification analyses were carried out using MNE-python (Gramfort et al., 2013) and scikit-learn (Pedregosa et al., 2011). Repetitions of individual words were averaged to increase signal-to-noise ratio (Grootswagers et al., 2017) and to avoid double-dipping in the cross-validation procedure. Time-resolved classification was achieved by inputting sequential 20 ms averages (5 time points) to a sliding classifier. We used a searchlight procedure across space and time to decode lexical information (in the lexical task only) and semantic information (in both tasks, separately). Lexical decoding was a pairwise classification procedure that aimed at distinguishing words (W) from pseudowords (P). Semantic information decoding was a multiclass problem (Abstract, A; unimodal Visual, V; Concrete-non-Visual, C), solved by separately classifying pairs of semantic classes (one-versus-one classification). Prior to each classification, classes were balanced by undersampling the class with the highest number of examples and by matching word duration between classes. Note that imbalance could occur only due to trial rejection. Whole-brain classification analysis was carried out at source-level. We used the Glasser atlas to parcellate the surface in 360 regions (Glasser et al., 2016). The vertices inside each parcellated region were used as features for the classification. Features were z-scored and given as input to a logistic regression algorithm with L1 penalty. Model performance was evaluated through a 13-fold stratified cross-validation scheme, to ensure a consistent number of examples of each class in the different folds. Each iteration yielded predictions of the left-out fold, evaluated using the ROC-AUC score (Fawcett 2006). The average score was then computed and assigned to the corresponding region. This procedure was repeated for all 360 regions at source level, resulting in 360 classification scores. The classification procedure was repeated at every time point, resulting in a time series of classification scores for each region. For the visual ROI analysis, we averaged the classification scores of the different parcels labelled as “early_visual” and “primary_visual” in the Glasser atlas (henceforth: “early visual”). In total 8 parcels were averaged, including both hemispheres.

In the lexical task, we further carried out a classification procedure of semantic classes focusing on abstract vs concrete (AV, AC) distinction. To maximise statistical power we used as features for the classifier all time points as well as all source space vertices, resulting in a time points x vertices feature vector for each trial. Classifier training and testing was then carried out as described above. This analysis resulted in a single classification score per cross-validation iteration, which was averaged to obtain one score per comparison and eventually averaged to obtain one semantic decoding score. This analysis was repeated considering as time points: all the time points between 0.4 s and 1 s; time points around the peak of semantic decoding in the semantic task ± 0.1 s. Semantic decoding peak was identified as the time point with highest numerical value within the significant cluster of the semantic task, collapsing the two groups together (cf Fig 3A).

**Figure 1.**
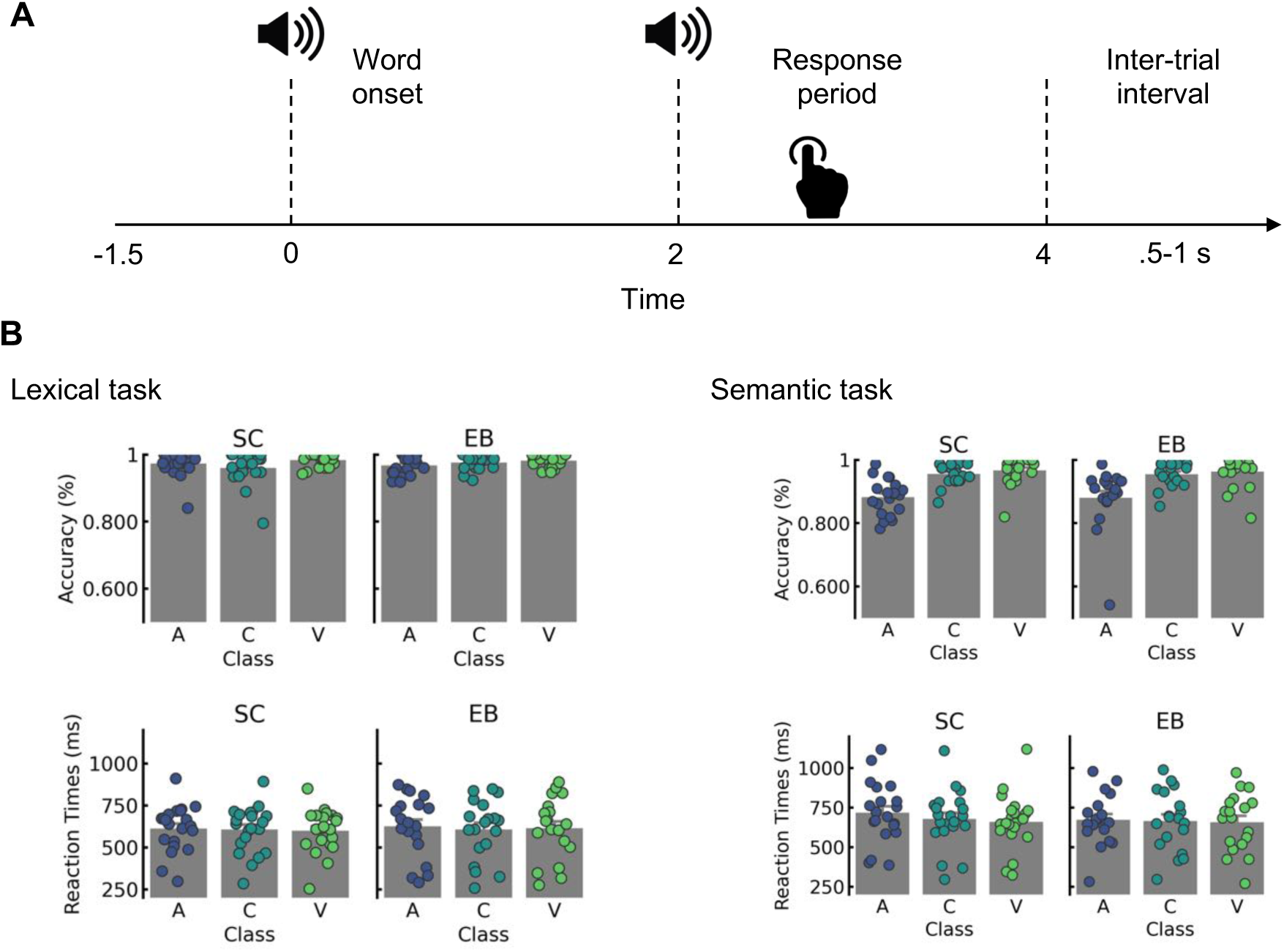
Experimental timeline and behavioral results A: Timeline of the experiment. Both sighted and blind participants entered the meg blindfolded, thus no visual stimulation was delivered. After an inter-trial interval of 1.5 s, participants were presented auditorily with a single word (t=0 word onset). After two seconds, a sound cue instructed them about the start of the response period. This period lasted 2 s, after which a jittered inter-trial period ended the trial. B: Behavioral results.Left: Lexical task: Top: response accuracy for each semantic class, sighted controls (SC) on the left and early blind (EB) on the right. Bottom: reaction times for each semantic class. Right: Semantic task: Same as lexical task. A: Abstract; C: Concrete non-visual; V: visual.

**Figure 2.**
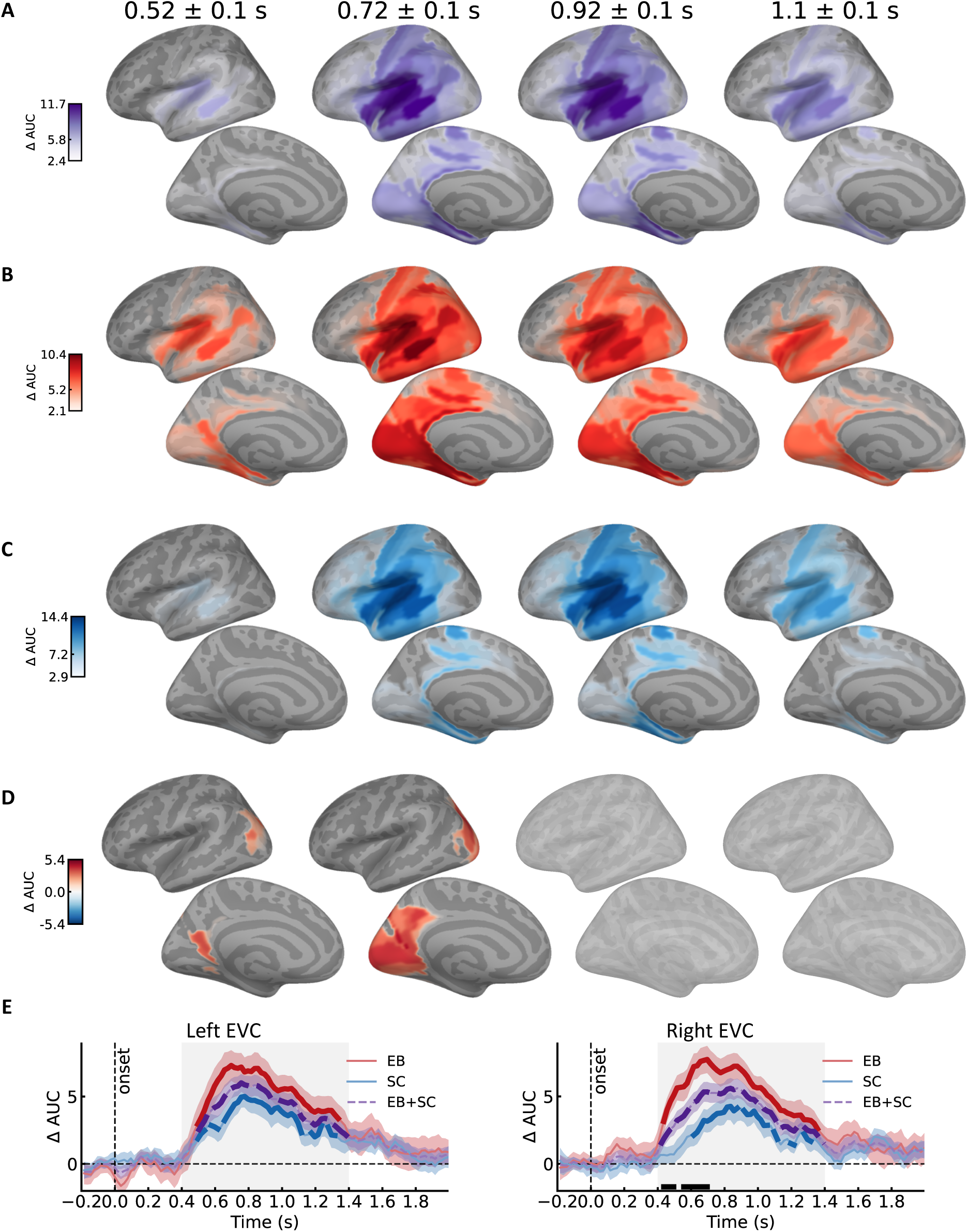
Lexical decoding A: Decoding performance when pooling groups together. Shown is the average decoding performance in four time windows, from left to right 0.52±0.1 s, 0.72±0.1 s, 0.92±0.1 s and 1.1±0.1 s. Brains are colored only in the significant parcels (p<0.05 cluster corrected over space and time), with darker color indicating higher decoding performance. Shown only left hemisphere, top row lateral view, bottom row medial view. See supplementary figure 1 for extended time course. B: Same as A, but showing results of the blind (EB) group. C: Same as A, but showing results of the sighted (SC) group. D: Same as A, but showing results of the contrast between groups. Red indicates EB>SC, blue indicates SC>EB. E: Decoding time course in early visual ROI timelocked to word onset. Left: left hemisphere ROI; right: right hemisphere ROI. Bold indicate time points of significant clusters within group. Black line indicate significant time points where EB>SC. Shaded area indicate time window of statistical testing (0.4 to 1.4 s).

**Figure 3a.**
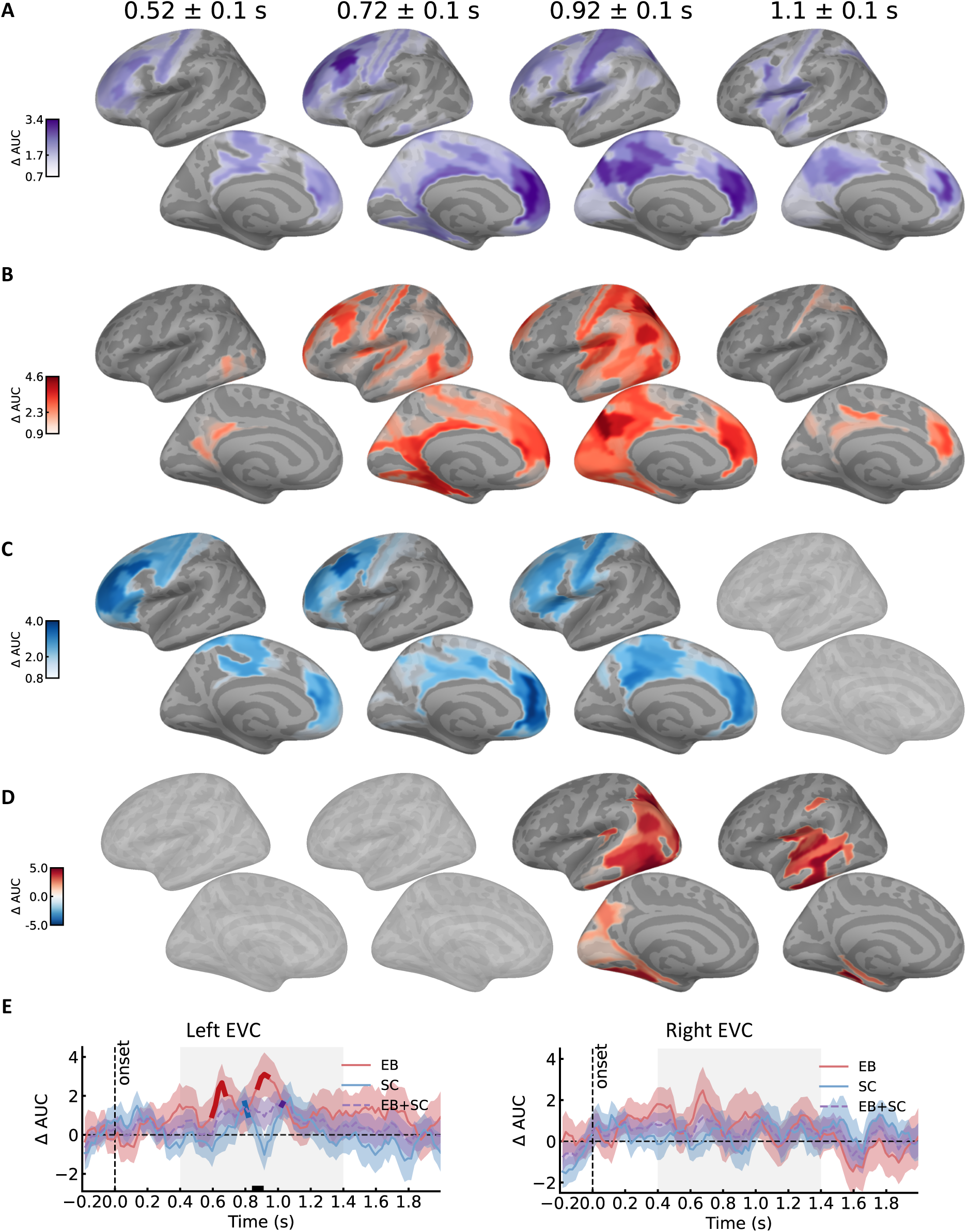
Semantic decoding in the semantic task A: Decoding performance when pooling groups together. Shown is the average decoding performance in four time windows, from left to right 0.52±0.1 s, 0.72±0.1 s, 0.92±0.1 s and 1.1±0.1 s. Brains are colored only in the significant parcels (p<0.05 cluster corrected over space and time), with darker color indicating higher decoding performance. Shown only left hemisphere, top row lateral view, bottom row medial view. For the full time course and right hemisphere see Supplementary Figure 2a. B: Same as A, but showing results of the blind (EB) group. C: Same as A, but showing results of the sighted (SC) group. D: Same as A, but showing results of the contrast between groups. Red indicates EB>SC, blue indicates SC>EB. E: Decoding time course in early visual ROI timelocked to word onset. Left: left hemisphere ROI; right: right hemisphere ROI. Bold indicate time points of significant clusters within group. Black line indicate significant time points where EB>SC. Shaded area indicate time window of statistical testing (0.4 to 1.4 s).

### Statistical analysis

For both EB and SC, we employed a cluster-based permutation approach (Maris & Oostenveld, 2007) to test whether the observed AUC score was significantly higher than chance level (0.5 in a binomial classifier) in a restricted 1s time window (i.e., 0.4 s to 1.4 s, Giari et al., 2020; Leonardelli et al., 2019). In brief, the cluster-permutation test first computes a univariate t-statistic at each data point. Data points surviving a one-sided (positive) uncorrected threshold of p<0.05 were then allowed to form clusters based on temporal and spatial adjacency. For the whole-brain analysis, temporal adjacency was trivially defined as the contiguous time points. Spatial adjacency was based on the distance between center of masses of each label (max distance 2 cm). The sum of the t-scores within the cluster was compared to a null distribution of clusters obtained from 10000 permutations. Based on the position of the observed cluster within this distribution a p-value for the cluster can be derived. A similar procedure was applied to test for differences in AUC between groups (i.e., EB vs SC), using a two-sided threshold at the cluster formation stage (cluster threshold p<0.05), given that we had no a-priori prediction about the direction of the effect. Finally, we extended this approach to the ROI analysis, with the only difference that neighbours were defined based on temporal adjacency only (no spatial information).

### Semantic information analysis

To assess the nature of semantic information processing, individual pairwise classification accuracies (AV, AC, and VC) sampled at each group’s peak decoding time (identified as the numerical maximum within the significant cluster) were analyzed using a repeated-measures Analysis of Variance (ANOVA). The model included Group (EB, SC) as the between-subjects factor and Class (AV, AC, VC) as the within-subject factor. A significant Group * Class interaction saw Class being investigated separately for each group using one-way ANOVAs. A significant Class effect was further investigated with post-hoc t-tests (one-sample, one-tail) conducted on each of the three classification accuracies, to assess above chance level performance. A Bonferroni correction was applied to the post-hoc tests, to control the family-wise error rate across multiple comparisons (Dunn, 1961). This analysis was repeated in three regions of interest: the early visual cortex, defined anatomically as described previously; the anterior cingulate/medial prefrontal cortex (ACC/mPFC) ROI and a posterior cingulate/precuneus ROI. These latter ROIs were functionally defined based on whole brain results. Specifically, we repeated the spatio-temporal whole brain cluster permutation test on the averaged classification accuracies collapsing across groups. For this analysis we selected as cluster forming threshold p<0.001 to improve reliability of selected data points. We then retained only clusters that contained at least 5 parcels, resulting in the two regions of interest.

## Results

Forty participants (19 early blind, EB; 21 sighted controls, SC) performed both a lexical decision task and a semantic decision task on the same set of spoken adjectives. By combining MEG, which affords millisecond temporal resolution and good spatial specificity, with multivariate classification analyses, we investigated lexical and semantic information processing in both groups. Above-chance classification was first assessed on a spatially unconstrained cortical surface (whole-brain analysis) and then within an a priori region of interest (ROI) targeting the sensory-deprived early visual cortex. Finally, we characterized group differences by directly comparing EB and SC decoding profiles in early visual cortex.

### Behavioral results

Tasks were designed to be easy and with minimal emphasis on execution speed; overall performance across groups was above 90% correct (LEX = 97.5±15.7%, SEM=93.4±23.8%) and reaction times were well within the 2s response interval (LEX = 609.5±269.4, SEM=673.9±308.4). We further investigated task-specific behavioral performance by entering both accuracy ratings and reaction times into separate repeated-measures ANOVAs with group (EB, SC) as a between-subject factor and class (A, C, V) as a within-subject factor. For the lexical decision task, analysis of response accuracy revealed a significant group × class interaction (F(2,76) = 3.701, *p* = 0.029). Follow-up one-way ANOVAs conducted separately for each group showed a main effect of class in EB (F(2,36) = 4.027, *p* = 0.026), and SC (F(2,40) = 5.751, *p* = 0.006). Post-hoc pairwise tests revealed higher accuracy for abstract vs visual words in EB (AV: t(18)=-2.935, p=0.008) while other contrasts showed no difference (AC: t(18)=-1.518, p=0.146; VC: t(18)=-1.215, p=0.239) whereas in SC we observed a lower accuracy for concrete non-visual words (AC: t(20)=2.281, p=0.033; VC: t(20)=-2.894, p=0.008) but no difference between abstract and visual (AV: t(20)=-1.517, p=0.144). Analysis on reaction times revealed a main effect of class (F(2,76) = 4.522, *p* = 0.013) with no main effect of group (F(1,38) = 0.03, *p* = 0.856) or interaction (F(2,76) = 1.371, *p* = 0.26). Post-hoc pairwise contrasts revealed a processing advantage for concrete words (V: 605.5±264 ms, C: 605.6±270.8 ms) over abstract words (A: 617.4±276.8 ms, i.e., concreteness effect; AV: t(39) = 2.523, *p* = 0.0158, AC: t(39) = 2.455, *p* = 0.0186; VC: t(39)=-0.196, p=0.845). This was confirmed in the semantic task behavioral performance. Analysis of accuracy rates showed a main effect of semantic class (F(2,76) = 22.251, *p* < .001) with no main effect of group (F(1,38) = 0.087, *p* = 0.769) or interaction (F(2,76) = 0.003, *p* = 0.996). Post-hoc pairwise contrasts revealed a processing advantage for concrete words (V: 96.5±18.4, C: 95.5±20.6) over abstract words (A: 88.2±32.3, i.e., concreteness effect; AV: t(39) = -5.157, *p* < .001, AC: t(39) = -4.754, *p* < .001; VC: t(39)=1.428, p=0.161). This was corroborated in the reaction times analysis, showing a main effect of semantic class (F(2,76) = 5.688, *p* = 0.004) with no main effect of group (F(1,38) = 0.128, *p* = 0.722) or interaction (F(2,76) = 1.777, *p* = 0.176). Post-hoc pairwise contrasts revealed a processing advantage for concrete words (V: 656.5±298.7 ms, C: 672.3±307.3 ms) on abstract words (A: 692.9±319 ms; AV: t(39) = 2.784, *p* = 0.008, AC: t(39) = 2.01, *p* = 0.05; VC: t(39)=-1.676, p=0.101).

Overall, task performance revealed a processing advantage for concrete over abstract words albeit task instructions putting little to no emphasis on execution speed.

### Lexical decoding

Speech recognition engages a hierarchically organized network of regions distributed along and around the Sylvian fissure, often referred to as the language network (e.g., Leonard et al., 2024). Within this system, regions such as superior and middle temporal cortices and the inferior frontal gyrus exhibit robust sensitivity to lexical information (e.g., Binder et al., 2009). Consistent with this framework, our data show robust classification of lexical information (words vs. pseudowords) during the lexical decision task in both groups (EB: p < 0.001, cluster corrected, Fig. 2B; SC: p < 0.001, cluster corrected, Fig. 2C). At the whole-brain level, classification peaks in EB and SC were located within canonical speech-processing regions, including primary auditory cortex and the posterior portions of the superior temporal gyrus, superior temporal sulcus, and middle temporal gyrus. Both groups displayed largely overlapping spatial and temporal profiles, with classification maxima between ∼0.6 and 0.8 s after stimulus onset (significant time windows: EB: 0.42–1.40 s; SC: 0.42–1.40 s).

Group differences were localized to posterior brain areas between 0.43 and 0.79 s (*p* = .029, cluster corrected), encompassing the cuneus, precuneus, inferior parietal cortex, and occipital cortex. Notably, higher classification accuracy in the posterior cortex of the blind emerged relatively early—approximately as early as the onset of significant decoding in classical language areas in both groups. In Figure 2E, we further characterize these posterior effects by constraining the analysis to “early visual” regions as defined by Glasser et al. (2016). Using a non-parametric cluster-based correction over time (Maris & Oostenveld, 2007), we found significantly above-chance lexical classification in early visual cortex in both EB (0.43–1.40 s, *p* < .001) and SC (0.57–1.11 s, *p* < .001, and 1.19–1.23 s, *p* = .024). Thus, early visual areas appear to respond to lexical stimulus properties in both groups. Crucially, however, the sensory-deprived early visual cortex of the blind showed significantly greater involvement during lexical processing than the non-deprived early visual cortex of sighted controls, with this effect being more pronounced in the right (EB > SC: 0.61–0.67 s, *p* = .012) than in the left hemisphere (p > 0.05).

### Semantic decoding

While convergent evidence suggests that visual areas in EB—and, as shown above, also in SC—are engaged in lexical processing, their role in semantic processing remains less well understood (Xu et al., 2023; Wang et al., 2020; Striem-Amit et al., 2018). In particular, it is unclear whether sensory-deprived visual cortices encode semantic information in a manner analogous to that of sighted individuals, and how task instructions shape top-down contributions to semantic retrieval.

To address these questions, we trained a classifier to discriminate among three well-defined semantic classes (Abstract, Concrete Visual, and Concrete non-Visual) and interpreted the average decoding accuracy as an index of overall semantic discriminability (Giari et al., 2020; Hebart et al., 2018). Crucially, in the lexical decision task, the instructions were orthogonal to semantic properties (all words were treated uniformly as lexical items), whereas in the semantic decision task participants explicitly attended to semantic attributes and classified words as abstract vs. concrete. Overall, this task manipulation exerted a strong influence on the observed brain activation and decoding profiles.

In the semantic task, semantic category information could be reliably decoded in both groups (EB: 0.576 -0.796 s, p = 0.03; 0.776 - 1.136 s, p=0.012) and SC (0.436-0.936 s, p=0.012). As illustrated in Figure 3A, this decoding engaged a widespread network closely matching the human semantic system described in prior meta-analyses and reviews (Binder et al., 2009; Lipkin et al., 2022). The network comprised posterior heteromodal association cortices (angular gyrus, middle temporal gyrus, fusiform gyrus, precuneus), subregions of heteromodal prefrontal cortex (including dorsal, ventromedial, and inferior–medial prefrontal cortex), and paralimbic regions such as the parahippocampal gyrus and posterior cingulate cortex.

Group differences (EB > SC: 0.836 - 1.136 s, p=0.022) were localized primarily to posterior and temporal cortices. More specifically, EB showed an initial (∼0.9 s) left-lateralized increase in decoding within inferior parietal cortex, cuneus, precuneus, and early visual cortex, followed by a later (∼1 s) enhancement in temporal regions including the anterior temporal lobe, ventral occipitotemporal cortex, middle and superior temporal gyri, supramarginal gyrus, and auditory cortex. Thus, during explicit semantic judgments, early visual areas contributed more strongly to semantic decoding in EB than in SC.

To further test this, we conducted planned ROI analyses in early visual cortices (EVC). These analyses confirmed a significant group difference in the left EVC, emerging around 0.85 s (0.85–0.89 s, *p* = .013). Importantly, semantic decoding in left EVC was not exclusive to the blind (0.596 - 0.676 s, p=0.014; 0.876 - 0.936 s, p = 0.028); in sighted participants, semantic information could also be decoded from the same ROI, albeit more transiently, with a peak around 0.8 s (0.796 - 0.816 s, p = 0.036).

What about semantic decoding during the orthogonal lexical decision task? At the whole-brain level, semantic category information yielded weak or non-existent above-chance classification. The only significant effect emerged in the left fronto-temporal cortex of EB participants (0.61– 0.99 s, *p* = .011), but no reliable group differences were observed (p > 0.05). Weaker semantic decoding under orthogonal task demands is broadly consistent with previous work reporting null or very limited semantic classification when semantics is task-irrelevant (Magnabosco & Hauk, 2024; Rahimi et al., 2022; Ruff et al., 2008).

However, behavioural evidence suggests that semantic information is nonetheless processed during lexical decision. Several studies have reported a robust concreteness effect in reaction times, with concrete words typically recognized faster than abstract words, even when the task is purely lexical (Connell & Lynott, 2012, 2016; Lynott & Connell, 2013). In line with this, a previous study from our lab showed a concreteness advantage for visually grounded, concrete non-visual vs. abstract words in both sighted and blind participants (Bottini et al., 2022). A behavioural result replicated in this study as well.

Motivated by these findings, we conducted an additional analysis focusing specifically on abstract vs. concrete distinctions (Abstract vs. Visual; Abstract vs. Concrete non-visual). To increase statistical power, we collapsed decoding across time and space (thus sacrificing temporal and spatial specificity). This analysis revealed significant above-chance decoding when considering both groups together (t(39)=1.857, p = 0.035, Supp. Fig. 3A), with no evidence for group differences (t(38)=-0.12, p=0.9). The effect was even stronger when restricting the analysis to a time window centred on the peak of semantic decoding (0.756 s) identified independently in the semantic decision task (t(39)=2.6, p = 0.006; Supp Fig. 3B).

**Figure 3b.**
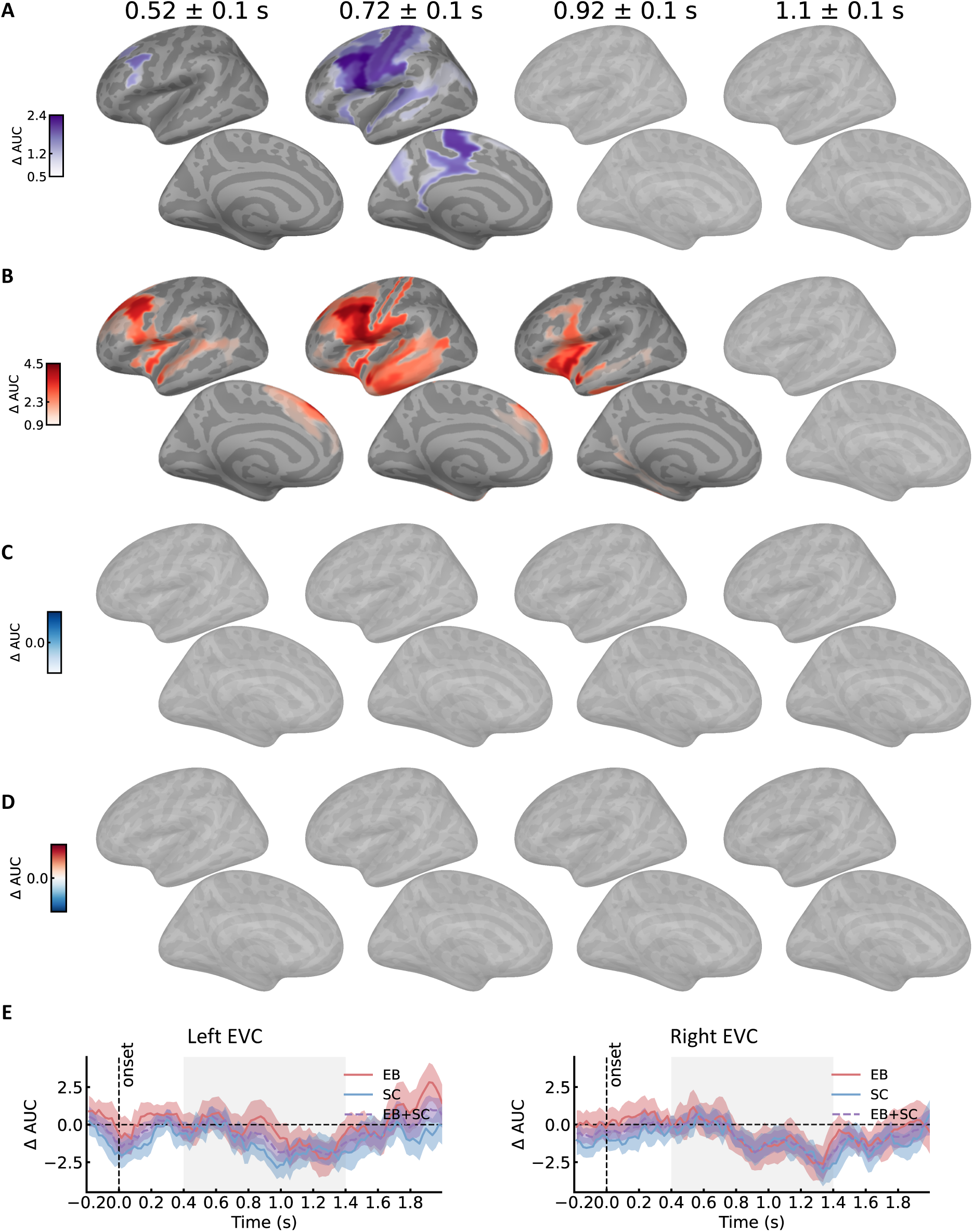
Semantic decoding in the lexical task A: Decoding performance when pooling groups together. Shown is the average decoding performance in four time windows, from left to right 0.52±0.1 s, 0.72±0.1 s, 0.92±0.1 s and 1.1±0.1 s. Brains are colored only in the significant parcels (p<0.05 cluster corrected over space and time), with darker color indicating higher decoding performance. Shown only left hemisphere, top row lateral view, bottom row medial view. For the full time course and right hemisphere see supplementary figure 2b. B: Same as A, but showing results of the blind (EB) group. C: Same as A, but showing results of the sighted (SC) group. D: Same as A, but showing results of the contrast between groups. Red indicates EB>SC, blue indicates SC>EB. E: Decoding time course in early visual ROI timelocked to word onset. Left: left hemisphere ROI; right: right hemisphere ROI. Shaded area indicate time window of statistical testing (0.4 to 1.4 s).

Crucially, none of these analyses showed significant semantic decoding within the EVC ROI (1s window: t(39)=-0.67, p = 0.747, Supp. Fig. 3C; decoding peak: t(39)=-0.24, p = 0.594; Supp. Fig. 3D). In sum, although not spatially or temporally localized, brain activity in both sighted and blind participants carried sufficient information to classify abstract vs. concrete words during lexical decision, paralleling previously reported reaction time concreteness effects in the same population (Bottini et al., 2022). Importantly, ROI analyses indicate that early visual cortex does not contribute detectably to this semantic discrimination when semantics is task-irrelevant.

### Semantic information

Thus far, we assessed semantic information processing as the aggregated pairwise classification accuracy across three contrasts: AV = Abstract vs. Visual; AC = Abstract vs. Concrete-non-Visual; and VC = Visual vs. Concrete-non-Visual. These analyses indicated that visual areas in both sighted controls (SC) and early blind (EB) participants are recruited during semantic processing, with a stronger overall involvement in the EB group. However, beyond this quantitative difference, semantic information might also be represented qualitatively differently in EVC across groups.

To further characterize the nature of semantic decoding in the semantic task in early visual cortex, we entered individual classification accuracies for AV, AC, and VC—sampled at the respective peak decoding times (0.876 s for EB; 0.796 s for SC)—into a repeated-measures ANOVA with group (EB, SC) as a between-subject factor and class (AV, AC, VC) as a within- subject factor. This analysis revealed a significant group × class interaction (F(2,76) = 3.64, *p* = .03). Follow-up one-way ANOVAs conducted separately for each group showed a significant main effect of class in EB (F(2,36) = 8.01, *p* = .001), but not in SC (F(2,40) = 0.304, *p* = .73).

In the EB group, planned Bonferroni-corrected post hoc comparisons indicated above-chance classification for AV (t(18) = 3.67, *p* = .003) and AC (t(18) = 4.00, *p* = .001), but not for VC (t(18) = 1.30, *p* = 1.00). This pattern suggests that, in early blind individuals, early visual areas are particularly sensitive to distinctions between abstract and sensory (visual or non-visual) concepts, whereas they do not reliably distinguish between different types of concrete sensory concepts (visual vs. non-visual).

Subsequently, we compared semantic decoding in EVC with two additional ROIs derived from the whole-brain semantic decoding results. Specifically, we created a thresholded mask from the pooled EB+SC group in the semantic task, restricted to the left hemisphere, and selected the two clusters showing the strongest and most spatially extended decoding in both groups (see methods): an anterior cingulate/medial prefrontal cortex (ACC/mPFC) ROI and a posterior cingulate/precuneus ROI.

In both ROIs, as expected, we observed significant semantic decoding in sighted and blind participants (posterior cingulate/precuneus: EB: 0.816 - 0.936 s, p = 0.001; 0.976 - 1.036 s, p = 0.013; SC: 0.976 - 1.096 s, p = 0.002; ACC/mPFC: EB: 0.656 - 0.736 s, p = 0.018; SC: 0.636 - 0.716 s, p = 0.024), with no reliable group differences in overall accuracy (p > 0.05). Two features of these profiles are particularly noteworthy. First, decoding accuracy peaked earlier in the anterior ACC/mPFC ROI (EB: 0.716 s; SC: 0.696 s) and later in the posterior cingulate/precuneus ROI (EB: 0.916 s; SC: 1.076 s), with the peak in the posterior cingulate closely aligned in time with the peak semantic effect in early visual areas (EB: 0.876 s; SC: 0.796 s). Second, a three-way anova with factors group (EB, SC) x roi (ACC/mPFC, posterior cingulate/precuneus) x class (AV, AC, VC) identified a significant roi x class interaction (F(1.70, 64.53)=5.33, p=0.01), indicating that the semantic profile differed across these regions. Indeed, aggregating across groups (justified by the absence of group x class interaction (F(1.96, 74.30)=0.45, p=0.637, see Supplementary Table 1 for the complete statistical report) in ACC/mPFC, all three pairwise contrasts reached above-chance levels (AV: t(39)=2.383, p=0.011; AC: t(39)=3.677, p<0.001; VC: t(39)=3.813, p<0.001), including the comparison between the two concrete classes (Visual vs Concrete-non-Visual). By contrast, the posterior cingulate/precuneus ROI exhibited a profile very similar to that observed in the EVC of blind participants (AV: t(39)=4.806, p<0.001; AC: t(39)=4.797, p<0.001; VC: t(39)=0.459, p=0.324) and in both groups with a strong task effect and selectively stronger decoding for abstract vs. concrete comparisons (AV and AC), but not for the concrete–concrete contrast (VC).

Interestingly, when considering all the three ROIs in Figure 4 in a three-way anova with factors group x roi x class, the 2-way interaction group x roi is not significant (F(1.80, 68.39)=0.39, p=0.655), showing that when the peak of the decoding effect is independently isolated in the two groups, the difference between groups in EVC is weakened (See supplementary table 2 for the complete statistical report).

**Figure 4.**
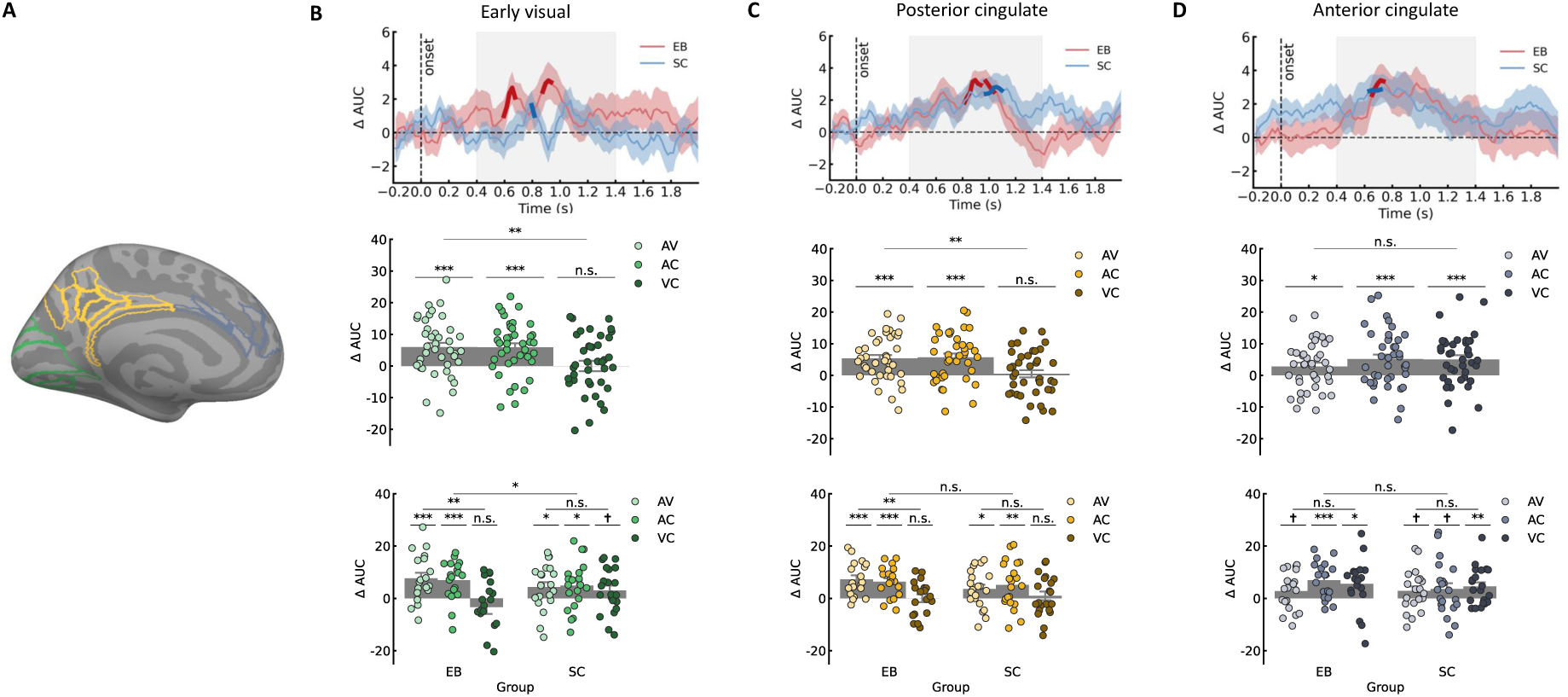
Semantic information and timecourse in selected ROIs: A: Visualization of the ROIs selected from the whole brain cluster permutation test plus early visual ROI. B: Top: Decoding time course in early visual ROI timelocked to word onset (same as figure 3a E). Decoding peaks at 0.876 s in EB and at 0.796 s in SC. Center: Decoding performance at the peak decoding time considering all EB and SC subjects together, separately for each pairwise comparison; Bottom: Decoding performance at the peak decoding time, split by group. C: Same as B but for posterior cingulate/precuneus ROI. Decoding peaks at 0.916 s in EB and at 1.076 s in SC. D: same as B but for anterior cingulate/medial prefrontal ROI. Decoding peaks at 0.716 s in EB and at 0.696 s in SC. (Abstract Visual: AV; Abstract Concrete non-visual: AC; Visual Concrete non-visual: VC) ***: p < 0.001; **: p < 0.01; *: p < 0.05; ✝: p< 0.1; n.s.: not significant

## Discussion

In both sighted and blind participants, the lexicality of a word could be decoded in a fronto-temporo-parietal network that highly overlaps with the human language system (Fedorenko et al., 2024; Binder et al., 1997; Lipkin et al., 2022), and the semantic category of a word (e.g., whether abstract, visual, or concrete but non visual) could be decoded in a partially distinct fronto-temporo-parietal network largely overlapping with the human semantic system (Binder et al., 2009). However, early blind participants consistently showed stronger decoding accuracy in early visual areas compared to sighted participants, both for lexicality and for semantic information. Remarkably, this was the case even though all stimuli were presented auditorily.

Such a finding is compatible with the idea that, in early blindness, early visual areas are reorganized to take up high-level cognitive processes such as lexical and, especially, semantic processing. However, although weaker than in the blind group, significant lexical and semantic decoding also emerged in the visual areas of sighted participants. This suggests a less radical view of brain plasticity, in which the occipital cortex of blind individuals carries out similar types of processes as the occipital cortex of sighted individuals. In what follows, we unpack our results in more detail to better characterize this hypothesis.

### Lexical processing in early visual cortex

Lexical decoding emerged in the early visual cortex (EVC) of blind participants, supporting previous evidence that deprived EVC is active during lexical processing in the blind (Bedny et al., 2011; Burton et al., 2003; Burton et al., 2002). For the first time, however, our MEG data allow us to access the time course of this activation. The involvement of EVC is relatively early: at the whole-brain level, a cluster in EVC emerges almost as early as clusters in fronto-temporal regions. This slight delay (see Supp. Fig. 1 for a higher temporal resolution visualization) is compatible with a cortico-cortical signal propagating from temporal language regions to EVC.

Interestingly, lexicality could also be decoded in the EVC of sighted people, although decoding accuracy was higher in the blind group. At the whole-brain level, the EVC cluster in sighted participants emerges more weakly and at later stages. However, this difference is drastically reduced in the ROI analysis and, in the left hemisphere, we did not find any significant difference between the decoding profiles of EVC in sighted and blind participants. The group difference in the right hemisphere might instead be driven by weaker language lateralization in early blind individuals, as reported in previous studies (Lane et al., 2017; Cheng, Silvano & Bedny, 2020; Tian et al., 2023; Dziegiel-Fivet & Jednorog, 2024).

Taken together, this pattern suggests that similar mechanisms are engaged in the EVC of sighted and blind participants during auditory lexical decision, with deprived EVC showing a stronger modulation. The role of the early visual cortex (EVC) in speech processing however remains to be fully determined. Previous studies in sighted participants have shown that the EVC exhibits stronger neural activation when listening to forward speech compared to backward speech (Seydell-Greenwald et al., 2023), responds to acoustic speech features related to intelligibility and vocal pitch (Venezia et al., 2021), and tracks the speech envelope in the absence of visual input (Bednaya et al., 2022; Hauswald et al., 2018; Van Ackeren et al., 2018). Here we show that this signal emerges relatively early (around 0.4 s), coinciding with activity in regions that showed the earliest statistically reliable differentiation between real words and pseudowords. These regions primarily include the auditory cortex and posterior temporal areas, such as the middle to posterior superior temporal sulcus (mid-post STS), which plays a key role in phonological processing and interfaces with the mental lexicon (Hickok & Poeppel, 2007). The timing of this effect aligns with the N400 component observed in event-related potential (ERP) studies, which consistently show significantly larger amplitudes for pseudowords than for real words in lexical processing tasks (see review by Lau et al., 2008). Converging evidence from fMRI research further demonstrates that auditory pseudowords elicit stronger activation in the superior temporal sulcus and gyrus than real words (Newman & Twieg, 2001; Kotz et al., 2002; Raettig & Kotz, 2008). These effects are commonly interpreted as reflecting attempts to interpret pseudowords given their phonological similarity to real words, or as increased phonological processing demands due to the absence of a lexical representation (Newman & Twieg, 2001; Kotz et al., 2002). Pseudowords may thus elicit a stronger prediction error that is propagated back to sensory cortices, within a predictive coding framework. Yet another possibility is that pseudowords induce greater effort or uncertainty, thereby recruiting a more extensive network that includes occipital regions.

Whatever the precise mechanisms, our data suggest that the processing in which EVC is involved during auditory lexical decision is not qualitatively different between sighted and blind participants. At the very least, there is no clear indication in our data that EVC in the blind supports a fundamentally different type of lexical computation. Consequently, these findings do not provide strong evidence for a radical functional reorganization of EVC in the lexical domain; instead, they support a view in which early visual cortex in blindness is more strongly integrated into a broadly similar lexical decision network.

### Semantic processing in early visual cortex

We show that semantic information can be decoded from early visual cortices (EVC) in blind individuals, in line with recent findings that the semantics of spoken words can be read out from occipital cortex in congenital blindness (Paczynska et al., 2025; Urbaniak et al., 2025; Noppeney et al., 2003; Abboud et al., 2019). Semantic decoding in blind EVC clearly emerges later than in classical nodes of the semantic network (Vignali et al., 2023), peaking around 0.9 s after word onset (Fig. 3a - Whole brain analysis). Importantly, this late peak is also the time window in which semantic decoding in EVC is significantly stronger in blind than in sighted participants (see both WB and ROI analysis).

This time course strongly suggests that a word’s semantic content is fully encoded and represented in canonical semantic regions (fronto-temporal and parietal heteromodal cortices) well before the signal reaches visual cortex. This raises reasonable doubts about whether EVC in the blind is a primary locus of semantic computation: it looks more like a recipient of top-down signals than a region where semantic categories are first constructed. However, this does not exclude the possibility that deprived EVC plays a functional role in semantic processing beyond simply mirroring the category structure computed elsewhere (for example by contributing to decision, prediction, or task control).

As with lexical processing, a similar effect emerged in sighted participants, especially in the left hemisphere and with a comparable late time course. Once again, this pattern suggests that we are witnessing a shared mechanism: early visual areas in both groups receive feedback from higher-level semantic regions, with blind EVC showing greater sensitivity to these top-down signals. The higher decoding observed in the blind may reflect increased responsiveness to feedback signals, possibly related to the supranormal metabolic activity at rest reported in early blind visual cortex or to an altered excitation–inhibition balance.

What type of signal is broadcast to EVC during the semantic task remains an open question. One possibility, already proposed in the literature, is that feedback to visual cortex selectively carries concrete semantic representations, with occipital cortex coding semantic dimensions tied to physical properties of referents (e.g., Urbaniak et al., 2025). However, if this was the case, one might expect similar semantic decoding in EVC also during the lexical decision task, where concrete and abstract words are present and semantic information is clearly processed in fronto-temporal regions in the blind. This was not the case: despite robust semantic coding upstream, semantic decoding in EVC was essentially absent when semantics was task-irrelevant.

A second, and in our view more likely, possibility is that EVC primarily receives task-related feedback carrying a low-dimensional, decision-relevant semantic signal. Our pattern in blind EVC—decoding of Abstract vs. Visual (AV) and Abstract vs. Concrete non-Visual (AC), but not of Visual vs. Concrete non-Visual (VC)—suggests that this signal approximates a semantic decision variable related to the degree of sensory grounding:

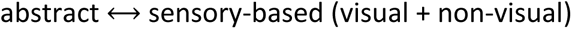

This account naturally explains why contrasts involving abstract vs. any concrete concept (AV and AC) are decodable, whereas the finer distinction between visual vs. non-visual concrete adjectives (VC) is not. Put differently, EVC does not appear to host a detailed, high-dimensional semantic feature map; instead, it seems to reflect a projection of a richer semantic state onto a behaviourally relevant axis—“how sensory-grounded/experiential vs. abstract is this word?”. Such a signal fits well within a predictive-coding framework, in which higher-order semantic regions send down action-relevant predictions about the expected level of sensory grounding for the current word, and these predictions are expressed in the activity patterns of early visual cortex.

This interpretation is supported by the decoding profile across semantic categories in blind EVC. In the blind group, EVC did not discriminate between visual and concrete non-visual words, consistent with the idea that its semantic response is dominated by the task-related abstract–concrete axis, rather than by fine-grained modality-specific content. Strikingly, this profile closely resembles the one observed in the posterior cingulate/precuneus of both sighted and blind participants, and contrasts with the pattern found in medial prefrontal cortex (mPFC). In mPFC, all three pairwise contrasts—including the concrete–concrete comparison—could be decoded, yielding a uniformly high decoding accuracy across semantic categories that was not strongly task-dependent. Moreover, semantic decoding in mPFC emerged earlier than in posterior cingulate and EVC.

Taken together, both the time course and the category-specific decoding profiles support a model in which EVC in the blind is recruited later than canonical semantic regions during semantic processing—consistent with a feedback modulation—and carries low-dimensional, task-relevant semantic information, similar to neighbouring posterior regions. In contrast, more anterior regions such as mPFC appear to support finer-grained, higher-dimensional semantic distinctions that are less tightly constrained by the current task.

Finally, it is worth noticing that, in the sighted, the EVC profile did not show such a clean abstract–concrete divide. This discrepancy may be due to the lower overall decoding accuracy found in the region for the sighted group, contributing to a less differentiated pattern across pairwise classifications. Indeed, also in the sighted EVC the classification between visual and non-visual words did not reach significance when considered alone. Another possibility is that visual simulations contributed to the decoding in the sighted. There is substantial evidence that, in sighted adults, listening to words can automatically trigger visual simulations of relevant properties (e.g., shape, colour, motion) in visual cortex (e.g., Fernandino et al., 2016). Critically, such visual simulations appear to be absent or strongly reduced in the blind (Bottini et al., 2020; Wang et al., 2020). Even relatively weak visual simulation in the sighted may make visual words decodable from concrete non-visual words in EVC, effectively adding a modality-specific layer on top of the shared task-related feedback signal. This added visual component could explain why the semantic profile of EVC in sighted participants diverges from that of blind participants, despite the overall similarity in timing and task dependence of semantic effects.

### Limitations

A limitation of the present study concerns the spatial resolution of MEG. Although source reconstruction allows us to estimate the origin of the signals at the cortical level, spatial precision is limited, and some of the spatial claims we make should be interpreted with caution. For example, the similar decoding profile observed in posterior cingulate/precuneus and in EVC raises the possibility that at least part of the effect attributed to EVC may reflect spillover or leakage from nearby posterior midline regions rather than activity originating strictly in early visual cortex.

We directly addressed this concern in supplementary analyses in which we restricted the ROI to BA17 (primary visual cortex) only, a more conservative and spatially circumscribed definition that should be less sensitive to contamination from precuneus or posterior cingulate. These analyses showed that semantic decoding in BA17 emerges when considering both groups together (p = 0.008 cluster corrected), with a single peak around ∼1 s post word onset and no significant difference between groups (p > 0.05 cluster corrected). Moreover, around this latency we observed significant decoding in the sighted group (p=0.047 cluster corrected) but not in the blind group (p > 0.05 cluster corrected), although, again, the direct comparison between groups did not reveal a significant difference (see Supplementary Fig. 4). Taken together, these BA17-restricted analyses are consistent with our main interpretation: early visual cortex, even under a strict anatomical definition, is engaged late and carries task-relevant semantic information, while group differences in this region are modest and do not suggest any radical functional reorganization due to early blindness.

### Conclusions

In conclusion, we found that the occipital cortices of both early blind and sighted individuals represent task-relevant lexical and semantic information. This finding highlights the preserved similarity of the sensory-deprived occipital cortex to that receiving sensory input, suggesting a more nuanced view of brain plasticity. Sensory-deprivation could thus lead to unmasking or upregulation of latent functions that are nevertheless already present.

## Data and Code Availability

The data supporting this study cannot be made publicly available because participants did not provide consent for their data to be shared. Code is available at https://github.com/giulianogiari/AbstractConcrete

## Author Contributions

LV, RB, YX, DC and OC designed the experiment. GG analysed the data with input from LV and RB. GG, LV and RB wrote the manuscript. All authors revised the manuscript.

## Declaration of Competing Interests

The authors declare no competing interests.

## Acknowledgements

We would like to thank our blind participants for participating in the study. This study was funded by a PRIN grant (Project number: 2015PCNJ5F_001) from the Italian Ministry of Education, University and Research (MIUR) awarded to Davide Crepaldi in collaboration with Olivier Collignon.

## Supplementary Results

**Supplementary Figure 1.**
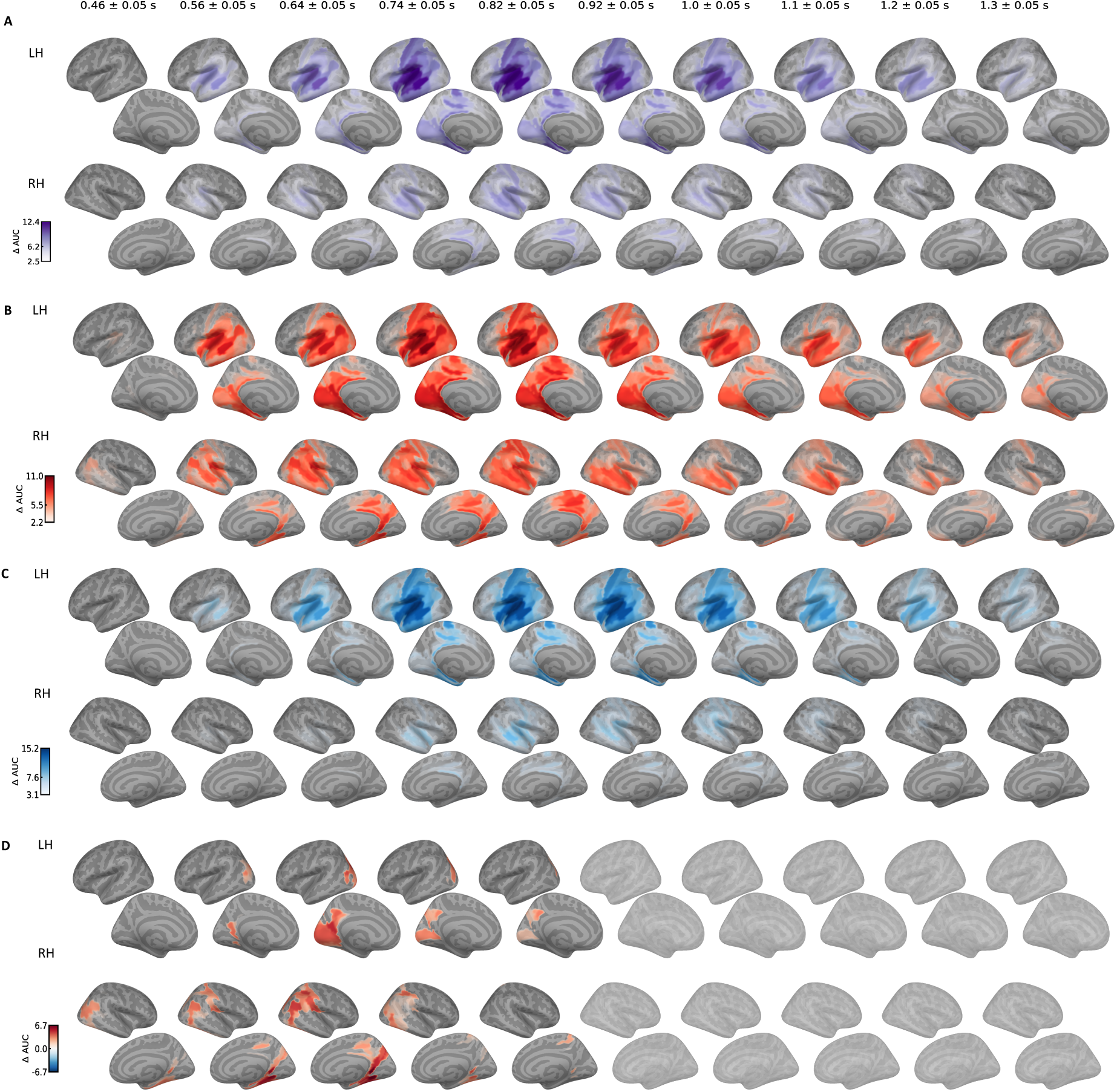
Lexical decoding in lexical task, shown both hemispheres with higher temporal resolution.

**Supplementary Figure 2a.**
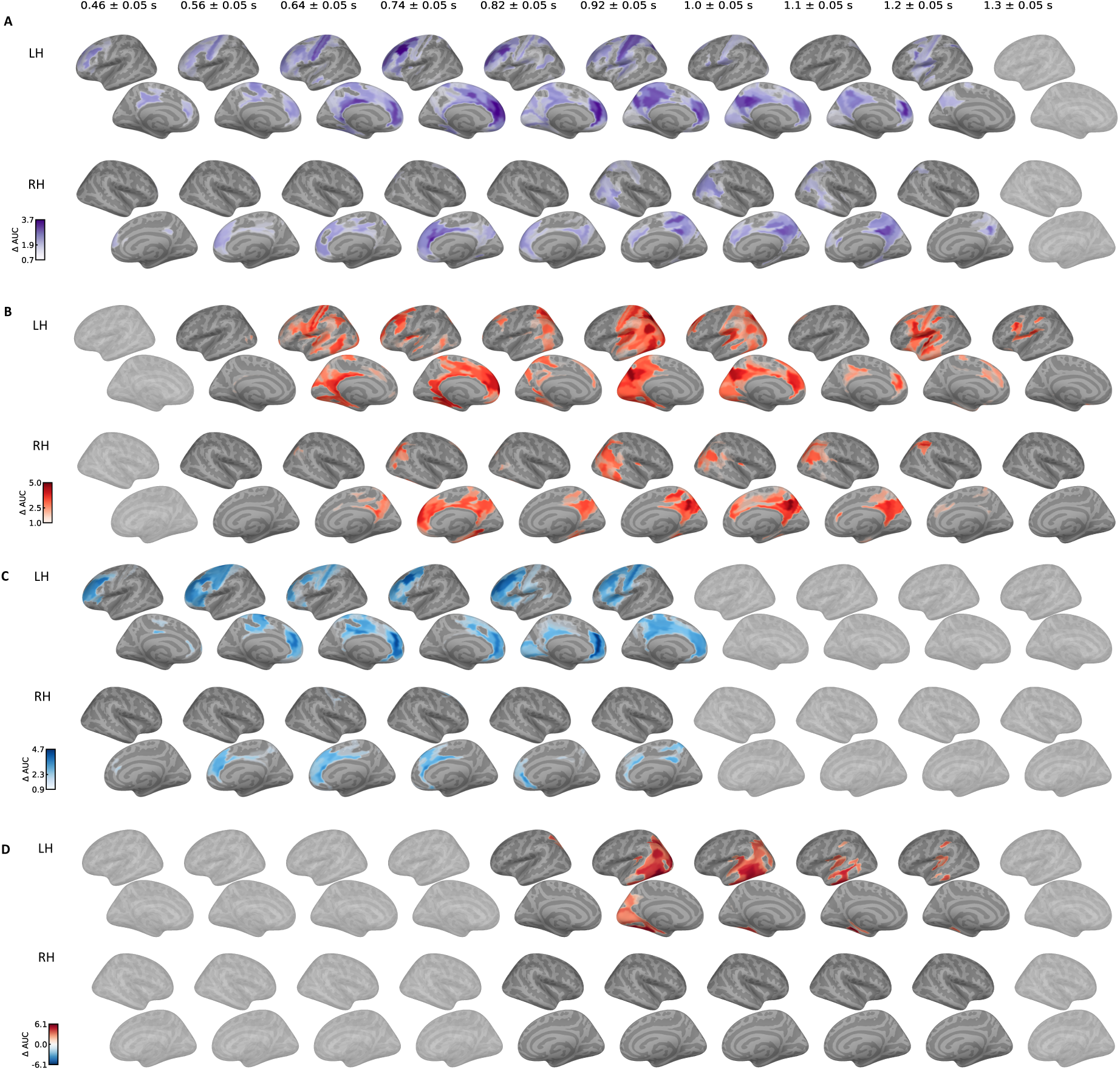
Semantic decoding in semantic task, shown both hemispheres with higher temporal resolution.

**Supplementary Figure 2b.**
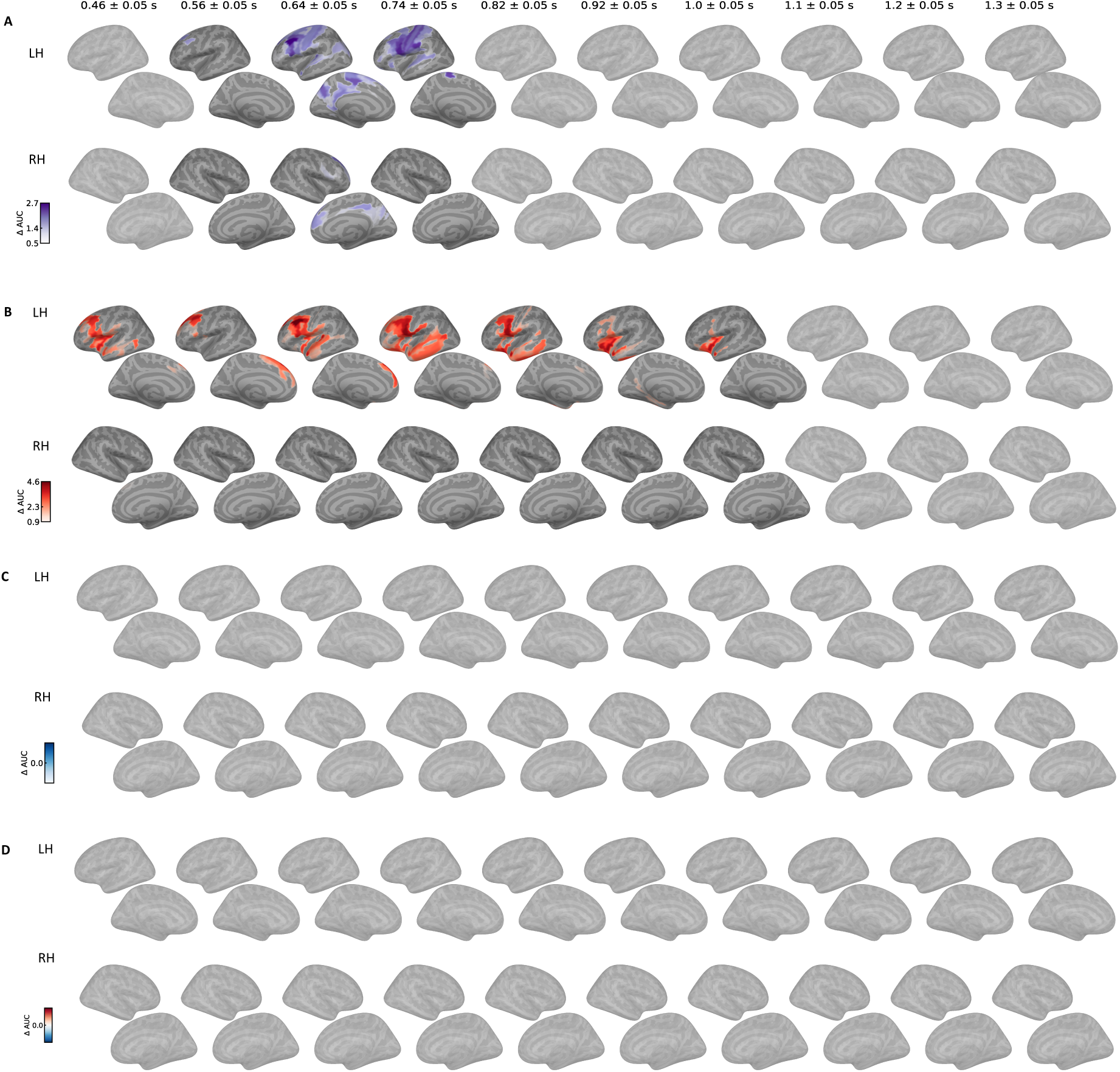
Semantic decoding in lexical task, shown both hemispheres with higher temporal resolution.

**Supplementary Figure 3.**
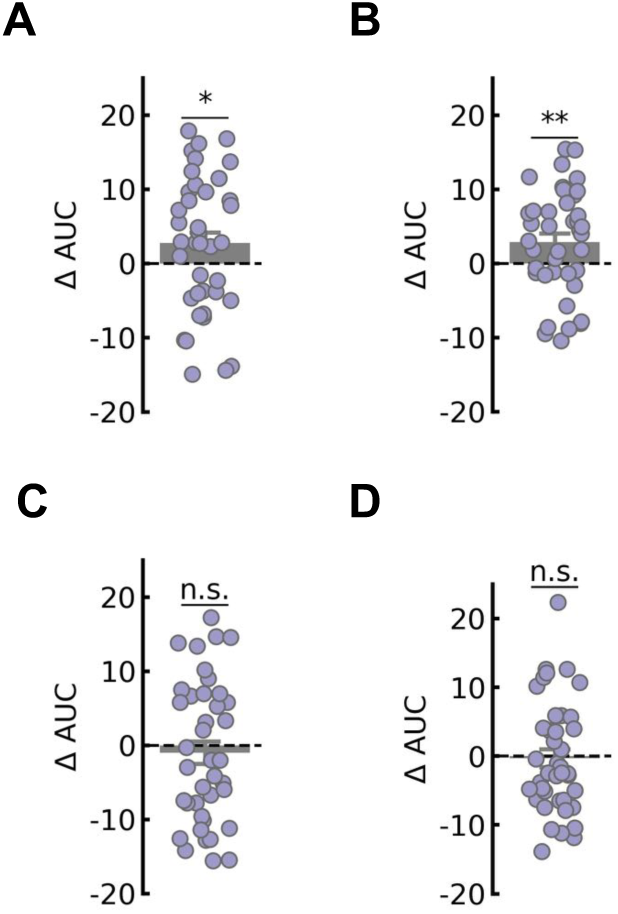
Semantic decoding in the lexical task, using all time points and vertices as features.

**Supplementary Figure 4.**
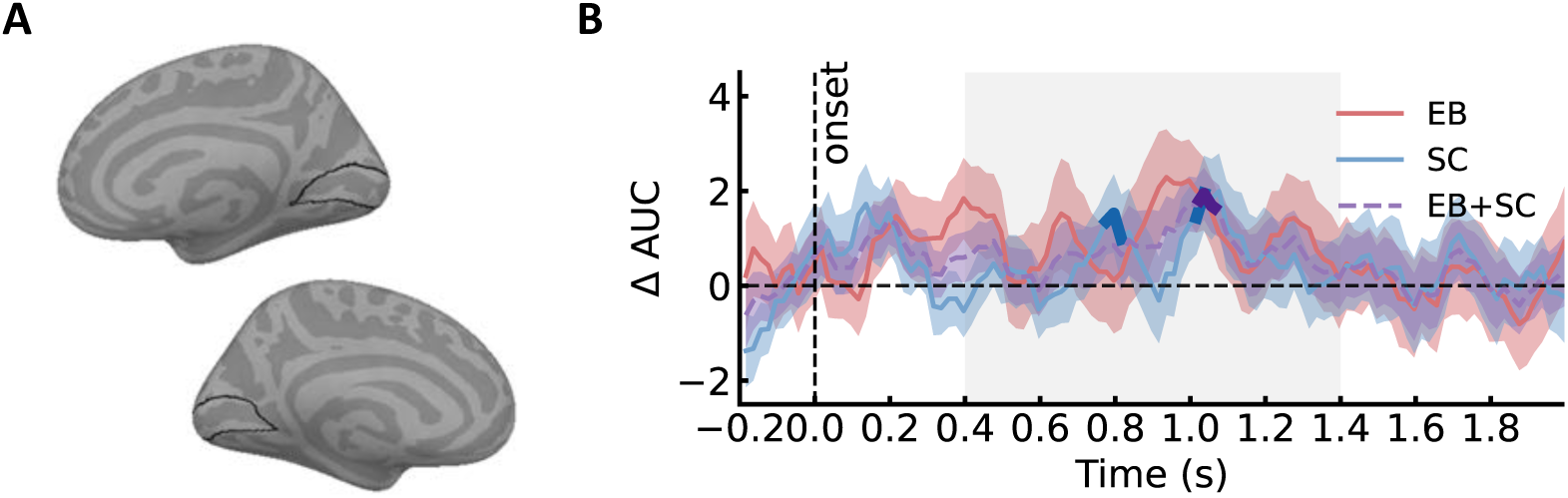
Semantic decoding in primary visual areas (BA17).

**Supplementary Table 1.** Three-way ANOVA with factors group (EB, SC) x roi (ACC/mPFC, precuneus) x class (AV, AC, VC).

| Effect | df | F | p |
| --- | --- | --- | --- |
| group | 1, 38 | 1.45 | 0.237 |
| class | 1.96, 74.30 | 2.3 | 0.109 |
| group x class | 1.96, 74.30 | 0.45 | 0.637 |
| roi | 1, 38 | 0.26 | 0.611 |
| group x roi | 1, 38 | 0 | 0.96 |
| class x roi | 1.70, 64.53 | 5.33 | 0.01 |
| group x class x roi | 1.70, 64.53 | 1.11 | 0.327 |

**Supplementary Table 2.** Three-way ANOVA with factors group (EB, SC) x roi (EVC, ACC/mPFC, precuneus) x class (AV, AC, VC).

| Effect | df | F | p |
| --- | --- | --- | --- |
| group | 1, 38 | 0.74 | 0.396 |
| class | 1.86, 70.80 | 6.73 | 0.003 |
| group x class | 1.86, 70.80 | 2.65 | 0.082 |
| roi | 1.80, 68.39 | 0.12 | 0.866 |
| group x roi | 1.80, 68.39 | 0.39 | 0.655 |
| class x roi | 3.41, 129.70 | 3.66 | 0.011 |
| group x class x roi | 3.41, 129.70 | 1.45 | 0.228 |

## Notes

### Competing Interest Statement

The authors have declared no competing interest.

